# Activation of bradykinin receptor B1 promotes desensitization of CXCR2 in neutrophils during severe sepsis and contributes to disease progression in mice

**DOI:** 10.1101/2024.04.19.590213

**Authors:** Raquel D N Arifa, Carolina B R Mascarenhas, Lívia C R Rossi, Maria Eduarda F Silva, Brenda Resende, Lívia D Tavares, Alessandra C Reis, Vanessa Pinho, Flavio A Amaral, Caio T Fagundes, Cristiano X Lima, Mauro M Teixeira, Daniele G Souza

## Abstract

Sepsis is one of the most common causes of death in intensive care units. The overproduction of proinflammatory mediators during severe sepsis leads to desensitization of CXCR2 on neutrophil, compromising their migration capacity. During early sepsis, kinins are released and bind to bradykinin 1 (BDKRB1) and bradykinin 2 (BDKRB2) receptors, however the involvement of these receptors in sepsis is not yet fully understood. This study demonstrated that the absence of BDKRB2 had no major effects compared to WT mice upon sepsis induction by CLP, suggesting that this receptor plays a minor role under these experimental conditions. In contrast, B1^-/-^ mice showed lower mortality and bacterial recovery compared to WT-CLP mice, which was associated with an increased influx of neutrophils into the peritoneal cavity of CLP-B1^−/−^ mice. WT-CLP mice exhibited increased expression of P110γ and decreased expression of CXCR2 in neutrophils, which was partially reversed in CLP-B1^−/−^ mice. Interestingly, local CXCL1 production was not affected by the absence of BDKRB1. In human neutrophils, LPS induced expression of BDKRB1, and antagonism of this receptor was associated with the restoration of neutrophil recruitment capacity upon stimulation with CXCL8. Furthermore, treatment with a BDKRB1 antagonist in combination with imipenem resulted in a significant improvement in mortality compared to animals treated with the antimicrobial agent alone. Our findings demonstrate that BDKRB1 plays an essential role in exacerbating the inflammatory response and CXCR2 desensitization in neutrophils during CLP-induced severe sepsis, highlighting BDKRB1 as a potential target for sepsis treatment.

**Importance:** Sepsis is a life-threatening organ dysfunction caused by a dysregulated host response to infection. Despite advances in understanding its pathophysiology, sepsis remains a leading cause of mortality in intensive care units nowadays. Here we found that B1 receptor contributes to neutrophil migration failure during severe sepsis. Inhibition of B1 improves neutrophil migration and bacterial clearance, making it a valuable therapeutic candidate for the treatment of sepsis. More importantly, treatment with a BDKRB1 antagonist in combination with imipenem resulted in a significant improvement in mortality compared to animals treated with the antimicrobial agent alone. These results highlight B1 as a potential treatment target for sepsis, offering improved modulation of the inflammatory response and synergy with antibiotics.

**Graphical Abstract:** BDKRB1 activation contributes to sepsis-induced hyperinflammation: (A) BDKRB1 activation contributes to sepsis-induced hyperinflammation: (A) BDKRB1 plays an essential role in the pathogenesis of sepsis, partly by mediating impaired neutrophil migration during the disease. It exerts its effects in myeloid cells by controlling the activation of P13Kγ and the expression of CXCR2. (B) BDKRB1 antagonist decreases cytokine production and increases neutrophil influx into the peritoneal cavity, resulting in a reduction in bacterial recovery, highlighting DALBK as a potential adjuvant treatment for sepsis

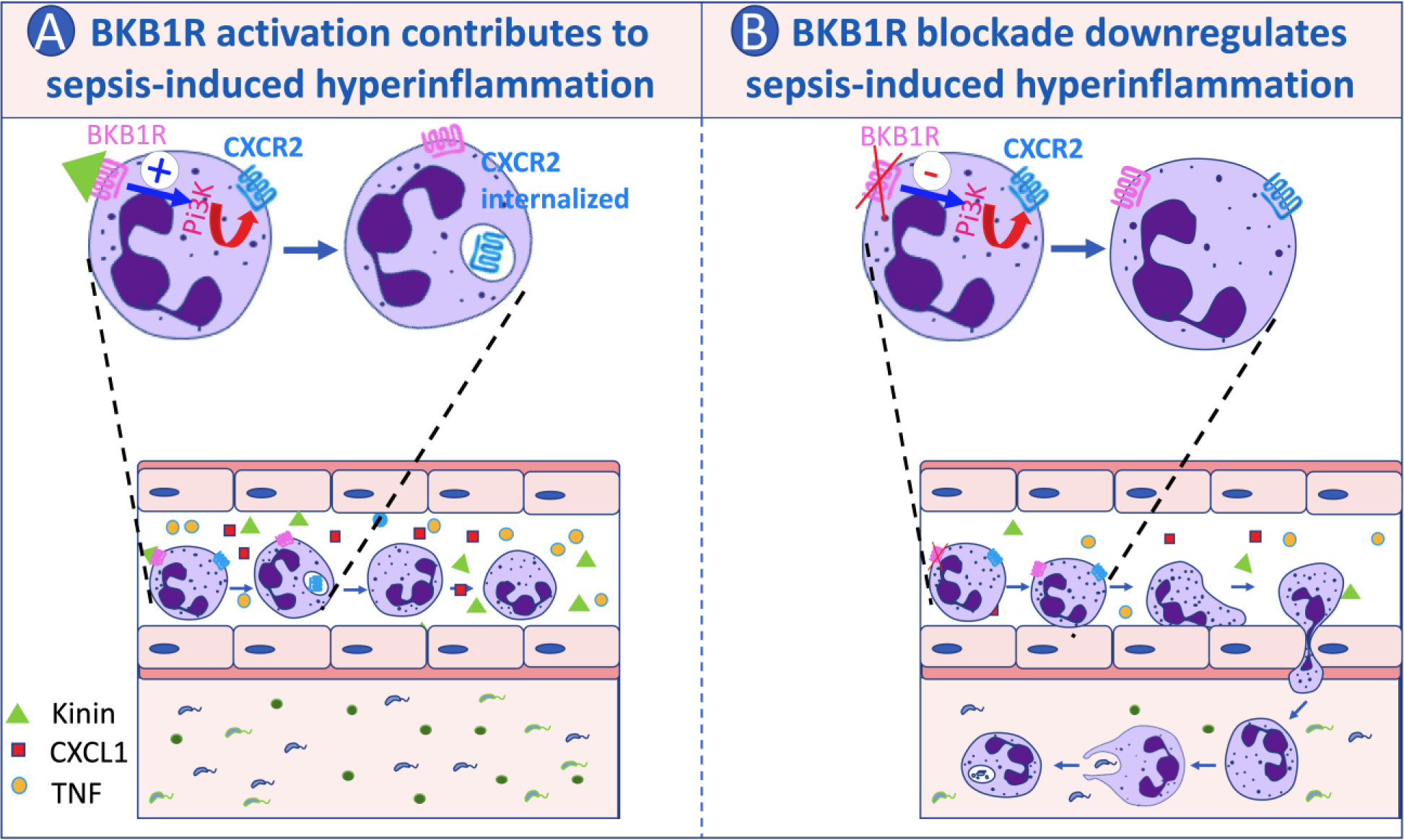

## 1. Introduction

Sepsis is described as potentially fatal organ failure due to an unbalanced host response to infection (1) and includes an earlier stage known as systemic inflammatory response syndrome (SIRS). In SIRS, elevated proinflammatory cytokines and widespread accumulation of inflammatory cells in vital organs can lead to organ failure, tissue damage and death. Neutrophils, the first line of defense against infection (2, 3), rely on the activation of CXCR2 receptors to effectively migrate to the site of infection (4). However, in sepsis, overproduction of proinflammatory mediators and increased activation of pattern recognition receptors trigger PI3Kγ, leading to phosphorylation of G protein-coupled receptor kinases (GRKs), that can desensitize CXCR2 on neutrophils (4). Reduced CXCR2 expression together with overexpression of CCR2 impairs neutrophil migration to the site of infection and favors their accumulation in the microvasculature of various organs, leading to multiple organ dysfunction syndrome (2, 3).

Bradykinin-related peptides are vasoactive molecules that play a role in various inflammatory conditions, including infection-related inflammation (5–7). These peptides are produced by kallikreins acting on kininogens and exert their effects via the activation of two different G protein-coupled receptors: BDKRB1 and BDKRB2 (8). BDKRB2 is constitutively expressed in various tissues and regulates several physiological processes (9). In contrast, BDKRB1 is only sparsely expressed in healthy tissue, but is upregulated by inflammatory mediators such as TNF and IL-1ß (8, 10). However, the role of BDKRB1 in the migration of leukocytes to sites of infection and its importance in sepsis are still poorly understood. The aim of this study was to investigate whether activation of the kinin receptor contributes to the inability of neutrophils to migrate to the site of infection in sepsis. The results show that BDKRB1 activation impedes CXCR2-mediated accumulation of neutrophils at the site of infection, resulting in impaired bacterial clearance, organ dysfunction and sepsis-related mortality.

## 2. Material and Methods

### 2.1. Animals

*Bdkrb1*^−/−^ (*B1*^−/−^) and *BdkrB2*^−/−^ (*B2*^−/−^) mice were generated as previously described (11). C57BL/6J-WT mice were purchased from the Biotério Central of the UFMG. 8–12-week-old mice were housed in separate cages under standard conditions and had free access to commercial food and water. The experiments were previously approved by the Animal Ethics Committee of the UFMG (Protocols 137/2012 and 136/2014).

### 2.2. Experimental model of severe polymicrobial sepsis induced by CLP

Mice underwent either severe sepsis by cecal ligation and perforation with an 18-gauge needle (CLP) or laparotomy only (sham control) as previously described (12). Survival of mice was monitored for 6 days after CLP. Mice were euthanized 6 hours after sepsis to collect peritoneal lavage, blood and lung samples.

### 2.3. Migration of leukocytes to the peritoneal cavity

Total leukocytes were counted using the Neubauer chamber. Differentiated counting was performed according to morphological characteristics. Quantification of the individual cell types was determined by the percentage of counted cells in relation to the total number of cells.

### 2.4. Bacterial count

Peritoneal lavage and blood samples were plated on Petri dishes containing Muller-Hinton agar medium (Difco Laboratories). After 24 hours of incubation at 37°C, the colonies were counted, and the results were expressed as the logarithm of colony forming units (CFU) per milliliter.

### 2.5. Myeloperoxidase assay

Neutrophil activity in the lung was determined by measuring myeloperoxidase (MPO) activity as previously described (13). Lung samples were frozen in liquid nitrogen and later processed to determine MPO activity. The change in optical density was measured at 450 nm using tetramethylbenzidine (Sigma-Aldrich). Results were expressed in relative units, comparing myeloperoxidase activity with that of casein-stimulated mouse peritoneal neutrophils processed in a similar manner.

### 2.6. Cytokine measurement

The concentrations of TNF, IL-6, IL-1β, IL-10, CXCL1 and CXCL2 were determined in the intraperitoneal lavage and in lungs; TNF and CXCL1 were determined in the serum. Cytokine concentrations were determined by ELISA using commercially available kits from R&D Systems (Minneapolis, MN, USA).

### 2.7. Histopathological analysis

The 5 µm thick lung sections were stained with hematoxylin and eosin. Lesions such as alveolar congestion, hemorrhage, neutrophil infiltration, aggregates in alveoli or vessel walls, and thickening of the interalveolar septum were identified and scored on a scale from 0 (absent) to 4 (severe) to standardize the assessment (14).

### 2.8. Isolation of neutrophils from blood and chemotaxis assay

Separation of human and mouse neutrophils (about 90% purity) was performed using the double-density histopaque method and quantified by trypan blue method. Chemotaxis assays were performed using a Modified Boyden’s chamber (Neuroprobe, Pleasanton, USA) and polycarbonate filters (4μm pores; Neuroprobe, Pleasanton, USA) as previously described (15). 1 x 10^6^ cells/mL of neutrophil were incubated with LPS (10µM) or vehicle for two hours. One side of the membrane received the CXCL8 (10 μM) while the other side was loaded with a neutrophil suspension. After 60 minutes at 37°C, the cells were quantified using Fast Panoptic Kit dye. The results represent the mean of 5 fields per membrane.

### 2.9. Chimera protocol

WT or *B1*^−/−^ mice were exposed to gamma radiation of 9 Gy with CO60 source in a Gammacell-220 irradiator. Bone marrow cells from WT or *B1*^−/−^ mice were transplanted intravenously into the irradiated recipient mice on the same day. The transplantation groups included WT-WT, *B1*^−/−^-*B1*^−/−^ *B1*^−/−^ WT, and WT-*B1*^−/−^ (donor-recipient, respectively). After cell injection, the mice received ciprofloxacin in their drinking water (70 mg/L) for 15 days. After a 10-day resting phase, mice were subjected to CLP sepsis.

### 2.10. Intravital Microscopy

Four hours after sepsis induction the rolling and adhesion of leukocytes to the mesenteric vascular endothelium was examined in according to established procedures (16). Leukocyte rolling was quantified as the number of rolling leukocytes per minute per vessel, and adhesion of leukocytes was confirmed when they remained on the vein endothelium for at least 30 minutes.

### 2.11. Immunofluorescence

The experiment was conducted two hours after surgery on mice. Neutrophils from WT CLP or sham and B1R^-/-^ CLP or sham mice were isolated and labeled with anti-GR1 and anti-P110γ antibodies to evaluate the fluorescence profile of P110γ in neutrophils. P110γ was quantified using a Mean Fluorescence Intensity (MFI). The MFI of P110γ was determined for 20 cells from each group of mice.

### 2.12. Western Blotting

The lung protein extracts were electrophoresed on a 12% polyacrylamide-SDS gel and transferred to nitrocellulose membranes according to established methods (8). The membranes were treated with specific primary antibodies (Santa Cruz Biotechnology, Santa Cruz, CA, USA), and then treated with the corresponding HRP-conjugated secondary antibody. The immunoreactive bands were visualized using an ECL detection system according to the manufacturer’s instructions (GE Healthcare, Piscataway, NJ, USA).

### 2.13. Flow cytometry

One hundred μL of blood from mice was incubated with anti-CD11b, anti-GR1, and anti-CXCR2 antibodies (R & D Systems). Neutrophils obtained from the blood of healthy individuals were stimulated with LPS (10µM) or vehicle for two hours and incubated with anti-CD14, anti-CD16, anti-B1R, anti-CXCR2, anti-PI3Kγ, and anti-rabbit antibodies (R & D Systems). Flow cytometric analysis was performed in FACSCanto II. The results were analyzed using the Flow Jo software.

### 2.14. Statistical analysis

Results were expressed as mean ± SEM for groups of 5 to 8 animals. Statistical analysis was performed using a nonparametric t-test or one-way analysis of variance (ANOVA ONE-WAY), followed by Tukey’s multiple comparison. The significance threshold was set at p < 0.05. Survival curves were presented as a percentage of live mice observed at 12-hour intervals over seven days. The Mantel-Cox-logrank test (chi-square, X2) was used for statistical analysis of survival curves, with differences considered significant at p < 0.05.

## 3. Results

### 3.1 B1, but not BKB2R, plays an important role in the pathogenesis of severe polymicrobial sepsis

In the severe sepsis model, 100% of WT or *B2*^-/-^ mice were dead 3 days after CLP. In contrast, *B1*^-/-^ mice began to die after 4 days and 50% were still alive on day 7 after CLP (Fig. 1A). Six hours after CLP, the bacterial load recovered in blood and peritoneal lavage was comparable in WT and *B2*^-/-^ mice, but about 2 log lower in *B1*^-/-^ mice (Fig. 1B). In septic WT and *B2r*^-/-^ mice, migration of leukocytes, especially neutrophils, into the peritoneal cavity was markedly restricted, whereas in *B1^-/-^*mice there was a significant influx of neutrophils (Fig. 1C, D). In WT mice, the inability of neutrophils to reach the site of infection was associated with reduced adherence to the endothelium, which was not the case in *B1*^-/-^ mice (Fig. 1E). Thus, activation of B1 impedes neutrophil migration and thus bacterial clearance in the CLP model, whereas BKB2R appears to be less important under these conditions. Therefore, *B2*^-/-^ mice were not used for further experiments.

**Figure 1:**
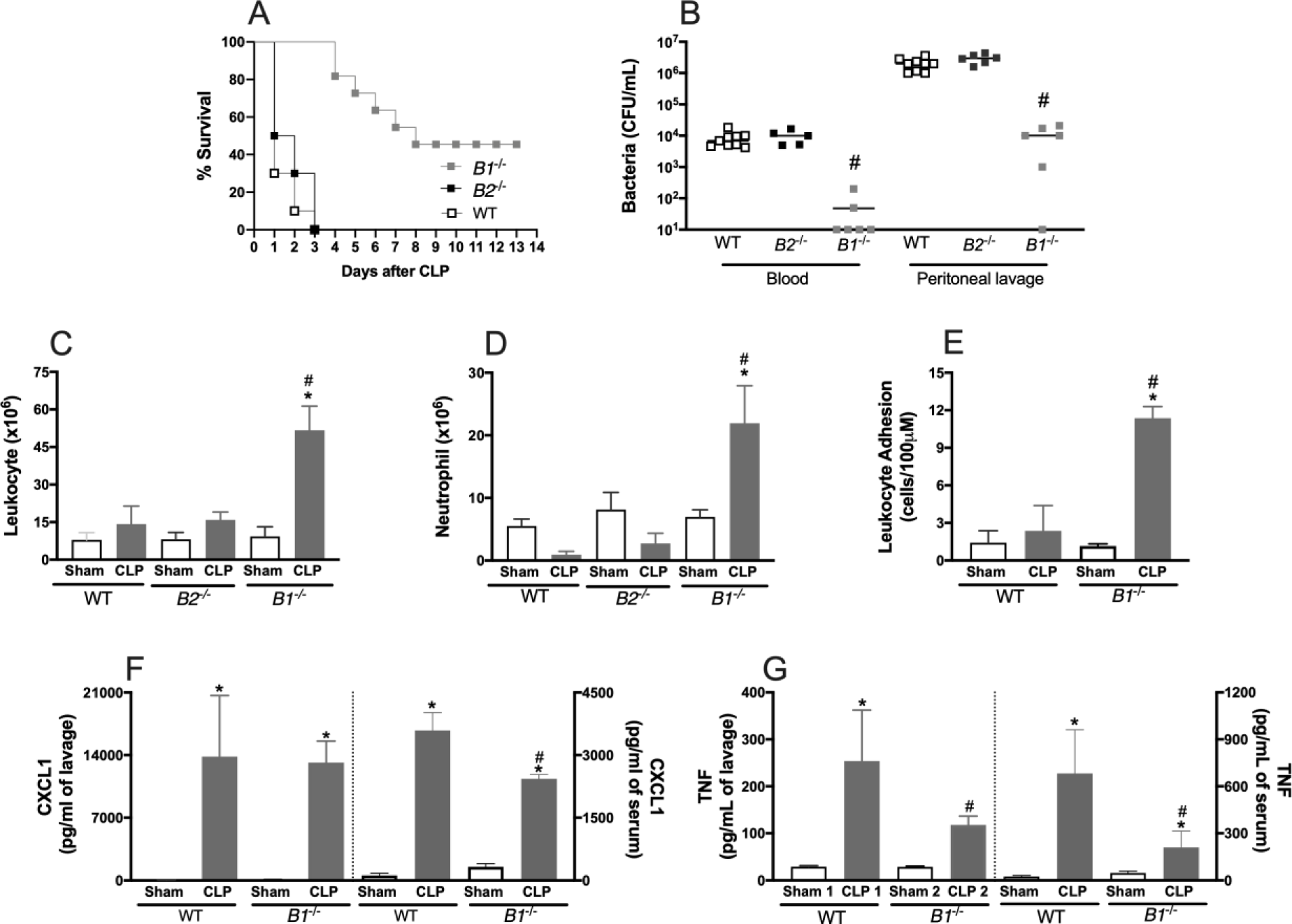
The kinin B1 receptor, but not the kinin B2 receptor, plays an important role in the pathogenesis of CLP sepsis. Wild-type mice (WT), bradykinin receptor 1-deficient mice (*B1*^-/-^), or bradykinin receptor 2 -deficient mice (*Br*^-/-^) were subjected to sham or CLP sepsis surgery. (A) The survival curve of mice subjected to CLP sepsis was followed for 14 days. Results are % of (A) survival, analyzed by the Log-rank (Mantel-Cox) test, 10 mice per group. Six hours after CLP sepsis, groups of mice were euthanized, blood was drawn and peritoneal lavage was performed. (B) Colony-forming units (CFU) in blood and peritoneal exudate, (C) total leukocytes, and (D) neutrophil recruitment in the peritoneal cavity were analyzed. (E) Leukocyte adhesion was analyzed 3 hours after CLP in the cremaster by intravital microscopy. Results are expressed as mean ± SEM of at least five animals per group. Six hours after CLP, groups of mice were euthanized, blood was drawn, and peritoneal lavage was performed. (F) TNF, and (G) CXCL1 concentration were quantified in the peritoneal lavage and serum. *, *p* < 0.05, compared to the sham group and ^#^, *p* < 0.05, compared to the wild-type mice subjected to CLP.

CLP led to increased concentrations of TNF and CXCL1 in the peritoneal lavage and serum (Fig. 1F-G) of WT mice. Except for TNF, *B1^-/-^*-CLP mice showed increased cytokine levels compared to *B1^-/-^*-sham mice in the peritoneal lavage (Fig. 1F-G). Interestingly, sepsis induced in *B1^-/-^*mice resulted in lower concentrations of TNF in serum and CXCL1 in both peritoneal lavage and serum compared to WT-CLP mice (Fig. 1F-G). CLP also led to increased CXCL2, IL-1ß, and IL-10 concentrations in the lavage fluid of both WT and *B1*^-/-^ mice, although there was no significant difference between the WT and *B1*^-/-^ mice subjected to CLP (Sup. Table 1). CLP also induced an increase in IL-6 production in the peritoneal lavage, but it is noteworthy that this cytokine concentration was reduced in *B1*^-/-^ mice (Sup. Table 1).

CLP induced significant lung injury characterized by severe capillary congestion, hemorrhage, infiltration of inflammatory cell, and alveolar edema (Fig. 2A) and increase of neutrophil influx into the lungs, as assessed by MPO activity (Fig. 2B). *B1^-/-^*mice showed improvement in all histopathologic parameters (Fig. 2A), resulting in a significant reduction in lung injury and MPO activity (Fig. 2B) compared to WT mice. The lungs of CLP-WT mice showed an intense production of cytokines, as shown by the increase in TNF (Fig. 2C) and CXCL1 (Fig. 2D). Indeed, CLP led to an increase of these cytokines in the lungs of *B1^-/-^* mice compared to the sham group. However, the levels of TNF (Fig. 2C) and CXCL1 (Fig. 2D) were decreased in CLP-*B1^-/-^* mice compared to CLP-WT mice. The production of CXCL2, IL-1ß, IL-6 and IL-10 in the lungs of WT and *B1*^-/-^ mice was increased. Interestingly, the concentrations of CXCL2 and IL-6 were reduced in *B1*^-/-^ mice compared to WT mice. (Sup. Table 1). These results suggest that B1 plays a central role in CLP-induced lung inflammation and injury.

**Figure 2:**
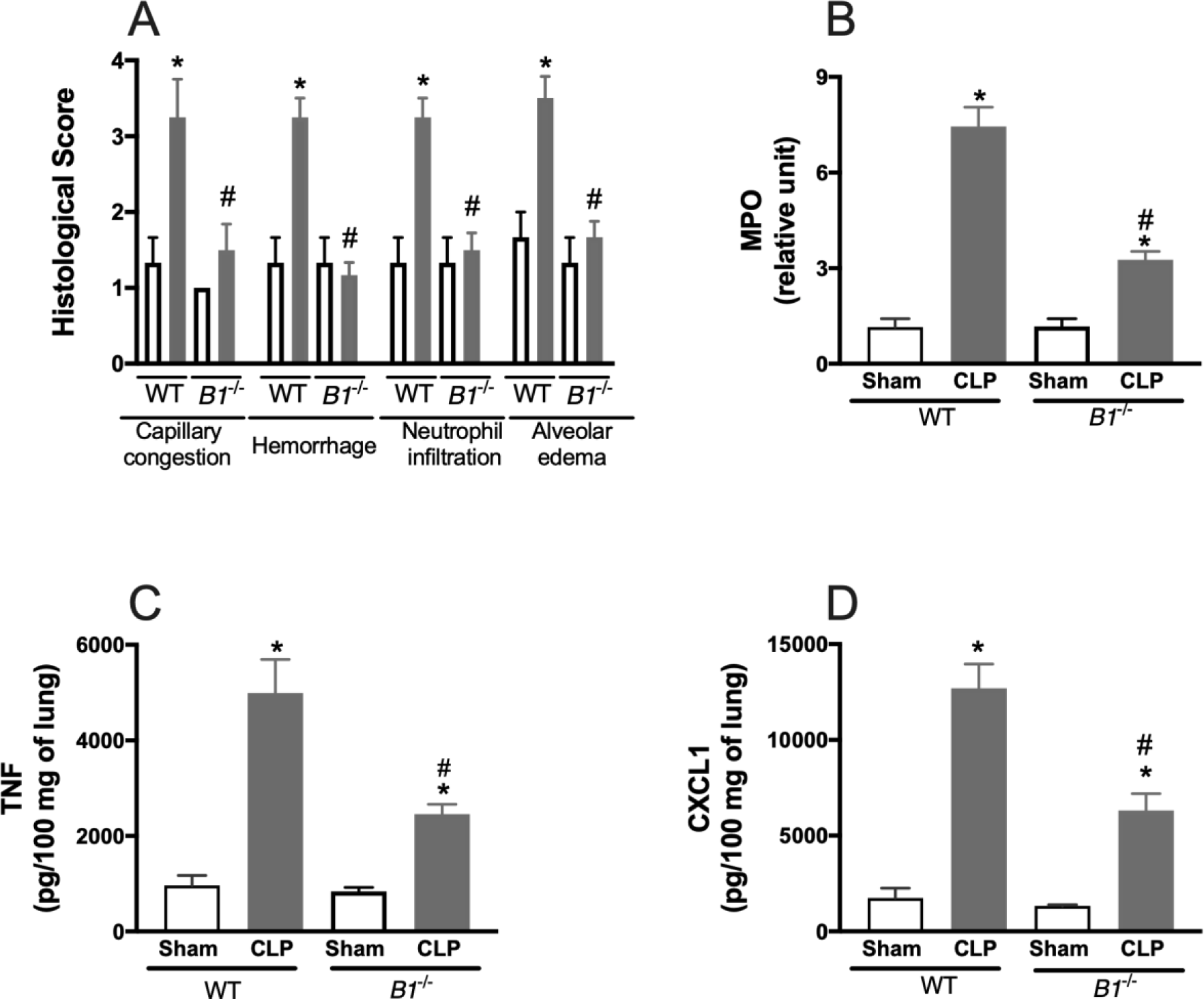
Absence of B1 prevents lung inflammation and injury. (A) Lung sections were stained with H&E for histopathologic analysis, tissue damage was quantified in three sections of each sample to assess capillary congestion, hemorrhage, neutrophil infiltration, and alveolar edema. (B) Neutrophil influx was determined by quantification of MPO activity. (C) TNF and (D) CXCL1 concentration in the lung were determined by ELISA. Results are expressed as mean ± SEM of at least five animals per group. *, *p* < 0.05, compared to the sham group and ^#^, *p* < 0.05, compared to wild-type mice subjected to CLP.

### 3.2 B1 in myeloid cells plays an essential role in exacerbating the inflammatory response triggered by severe sepsis

To investigate the role of the B1 receptor in myeloid cells in CLP sepsis, *B1*^−/−^ and WT mice were irradiated and subsequently transplanted with bone marrow-derived cells (BMDC) from *B1*^−/−^or WT mice. WT mice receiving *B1*^−/−^ BMDC (*B1*^−/−^→ WT) exhibited increased peritoneal leukocyte (Fig.3A) and neutrophil (Fig. 3B) counts and reduced local (Fig. 3C) and systemic (Fig. 3D) bacterial recovery after CLP compared with *B1*^−/−^ mice transplanted with WT BMDC (WT → *B1*^−/−^). Furthermore, *B1*^−/−^ → WT mice exhibited similar cytokine levels and neutrophil influx into the lungs as *B1*^−/−^ mice receiving *Br1*^-/-^ BMDC (*B1*^−/−^ → *B1*^−/−^) (Fig. 3B, E-K). In contrast, *B1*^−/−^ mice receiving BMDC from WT (WT → *B1*^−/−^) showed the same phenotype as WT mice receiving WT BMDC (WT → WT): no neutrophil migration (Fig. 3B and 3K) and consequently a high bacterial burden (Fig. 3C-D) after CLP. In addition, WT → *B1*^−/−^ mice had higher concentrations of TNF and CXCL1 in the peritoneal cavity, blood and lungs compared to WT → *B1*^−/−^ mice (Fig. 3E-J). These results emphasize that the role of B1 in sepsis depends on its presence in myeloid cells, which are critical for the migration of neutrophils to the site of infection and for the regulation of the inflammatory response triggered by CLP.

**Figure 3:**
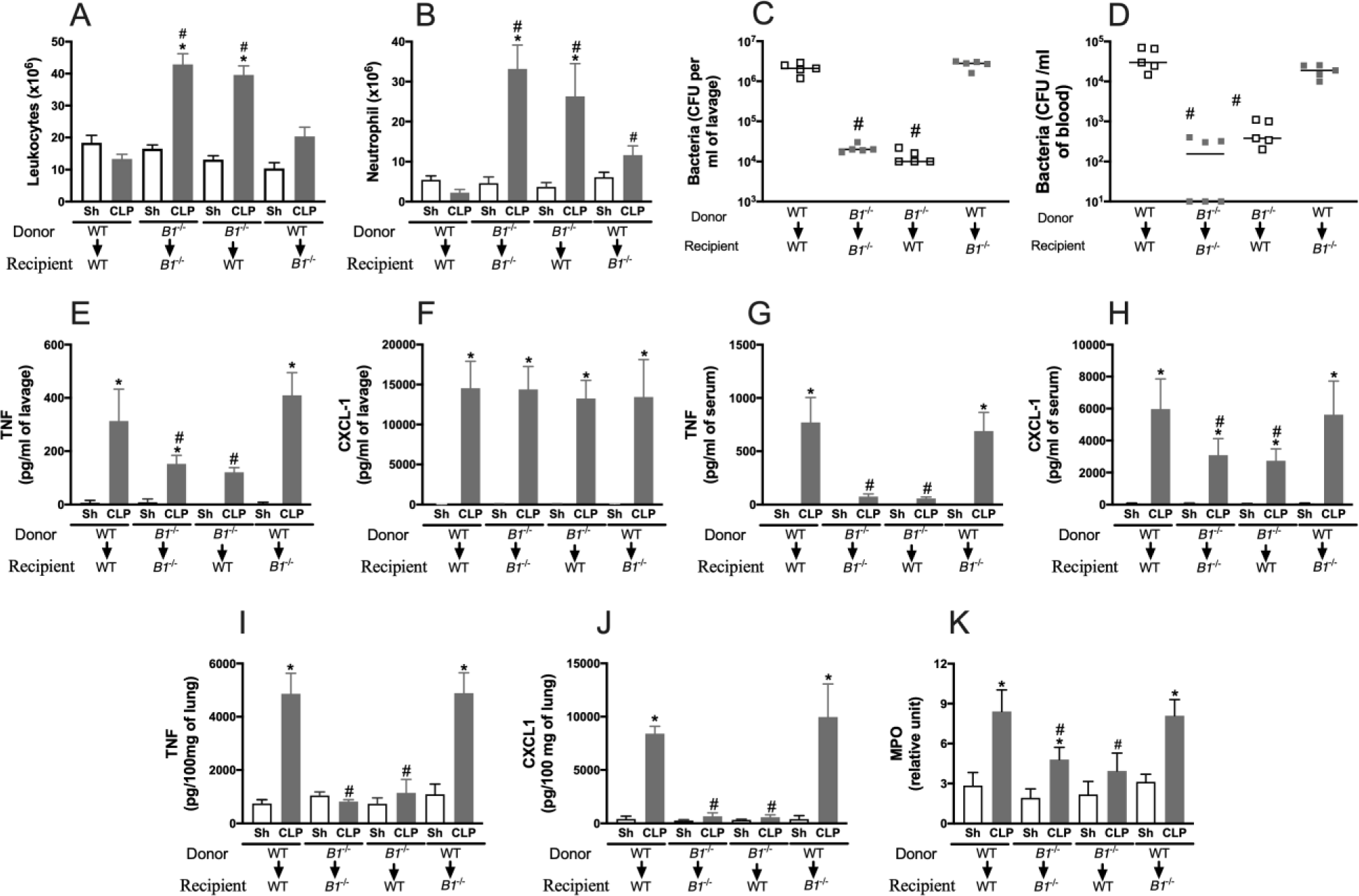
B1 in myeloid cells plays an essential role in exacerbating the inflammatory response induced by CLP. Chimeric mice were generated by the adoptive transfer of 10^6^ BM cells from donor to recipient mice. Four groups of mice were generated: cells from WT mice to WT mice, cells from *B1*^-/-^ mice to *B1*^-/-^ mice, cells from *B1*^-/-^ mice to WT mice, and cells from WT mice to *B1*^-/-^ mice. Each group was divided into two groups that underwent to sham or CLP surgery. Six hours after CLP, the groups of mice were euthanized, blood and lungs were collected, and peritoneal lavage was performed. (A) Total leukocytes, (B) neutrophil influx into the peritoneal cavity. The bacterial load in (C) blood and (D) peritoneal exudate was determined by plating and CFU counting. The concentrations of (E) TNF and (F) CXCL1 were determined in peritoneal lavage, blood (G-H) and lung (I-J). (K) MPO activity was determined in the lung. Results are expressed as mean ± SEM of at least five animals per group. *, *p* < 0.05, compared to the sham group and ^#^, *p* < 0.05, compared to wild-type mice subjected to CLP.

### 3.3 B1 activation induces increased Pi3Kγ expression and desensitization of CXCR2 on neutrophils

CLP induced a decrease in both the percentage (Fig. 4A) and number of neutrophils (Fig. 4B) expressing CXCR2 in WT mice, accompanied by a decrease in the mean fluorescence intensity (MFI) of CXCR2 on these cells (Fig. 4C). Furthermore, CLP-induced sepsis resulted in upregulation of P110γ, the catalytic subunit of PI3Kγ in WT mice compared to WT-sham mice (Fig. 4D and 4E). The absence of B1 resulted in a reversal of CXCR2 desensitization and P110γ activation, which were very similar to the profiles observed in sham-operated mice. Collectively, these results suggest that B1 disrupts neutrophil migration by desensitizing CXCR2 in response to PI3Kγ activation.

**Figure 4:**
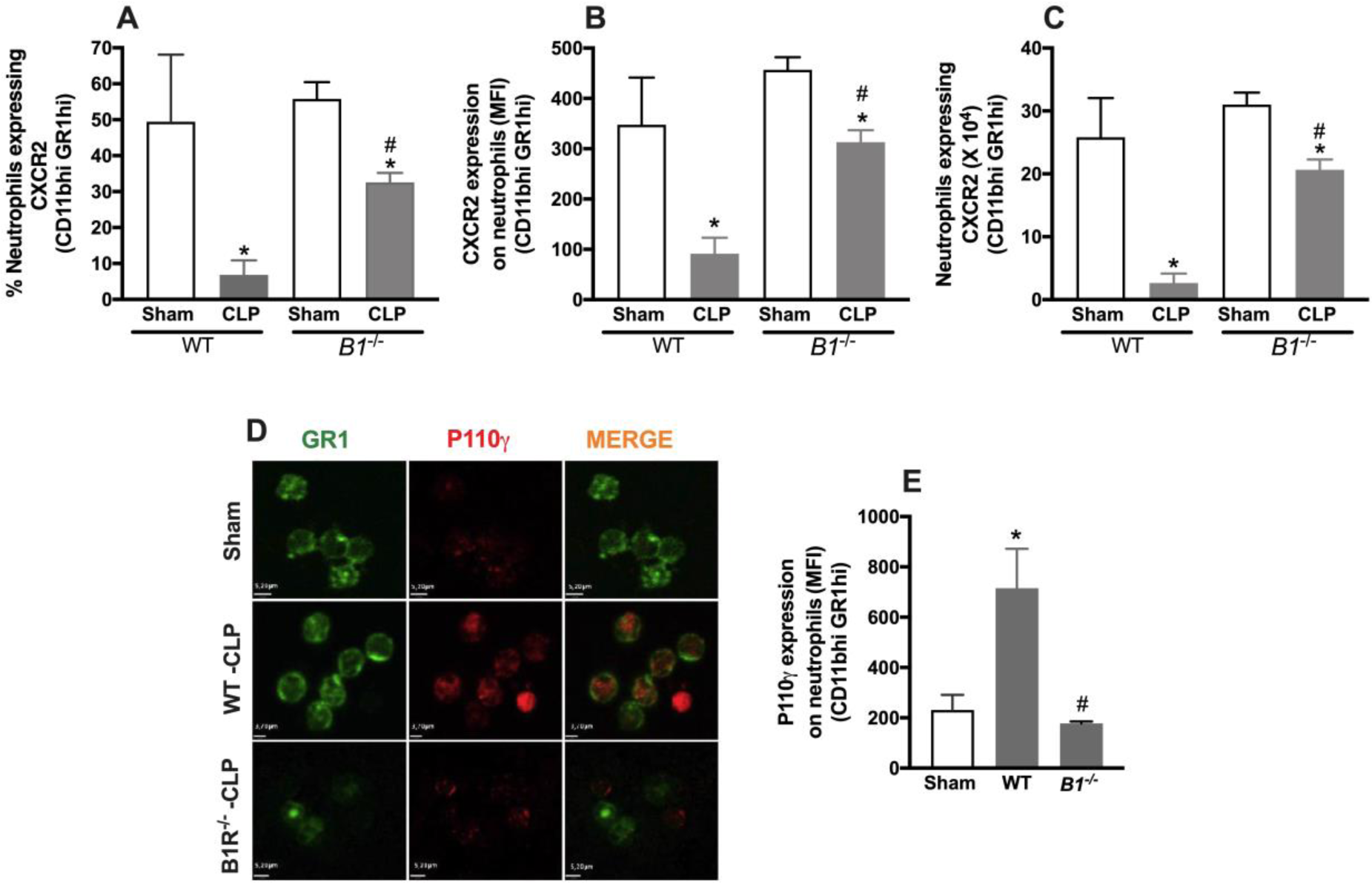
B1 activation induces Pi3Kγ activation and desensitization of CXCR2 receptor in circulating neutrophils after CLP-induced sepsis. (A) Percentage of neutrophils positive for CXCR2 receptor labeling, (B) Number of circulating neutrophils positive for CXCR2 and (C) Mean Fluorescence Intensity in neutrophils of septic mice. Results are expressed as mean ± EPM of five animals per group. * p<0.05 for the sham group from the same strain (controls), # p<0.05 related to the WT sham group. (D) Representative immunofluorescence image showing the fluorescence profile of neutrophils (labeled with anti-Gr1 antibody) and P110γ (labeled with anti-P110γ antibody), as well as the image overlay (merge). Two hours post-surgery, circulating neutrophils of WT CLP or sham and B1R^-/-^ CLP or sham mice were evaluated. (E) p110γ was quantified as the Mean Fluorescence Intensity (MFI) of 20 cells from each group. * p<0.05 related to the Sham group and # p<0.05 related to the WT CLP group

Human neutrophils stimulated with LPS showed an increased percentage of neutrophils expressing B1 (Fig. 5A) and a higher MFI (Fig. 5B) of this receptor. In addition, pre-incubation with LPS led to a decrease in the percentage of neutrophils expressing CXCR2 (Fig. 5C), decreased MFI of CXCR2 (Fig. 5D) and increased P110γ expression (Fig. 5E). Pre-incubation with LPS reduced neutrophil transmigration induced by CXCL8. Interestingly, incubation of neutrophils with the B1 agonist, DABK, or the B1 antagonist, DALBK, had no effect on CXCL8-induced chemotaxis (Fig. 5F). However, treatment with DALBK reversed the LPS-induced reduction in neutrophil migration. However, DABK did not affect the inability of neutrophils pre-stimulated with LPS to recruit towards the CXCL-8 gradient. Thus, inhibition of B1 in human neutrophils inhibits their ability to migrate under sepsis conditions.

**Figure 5:**
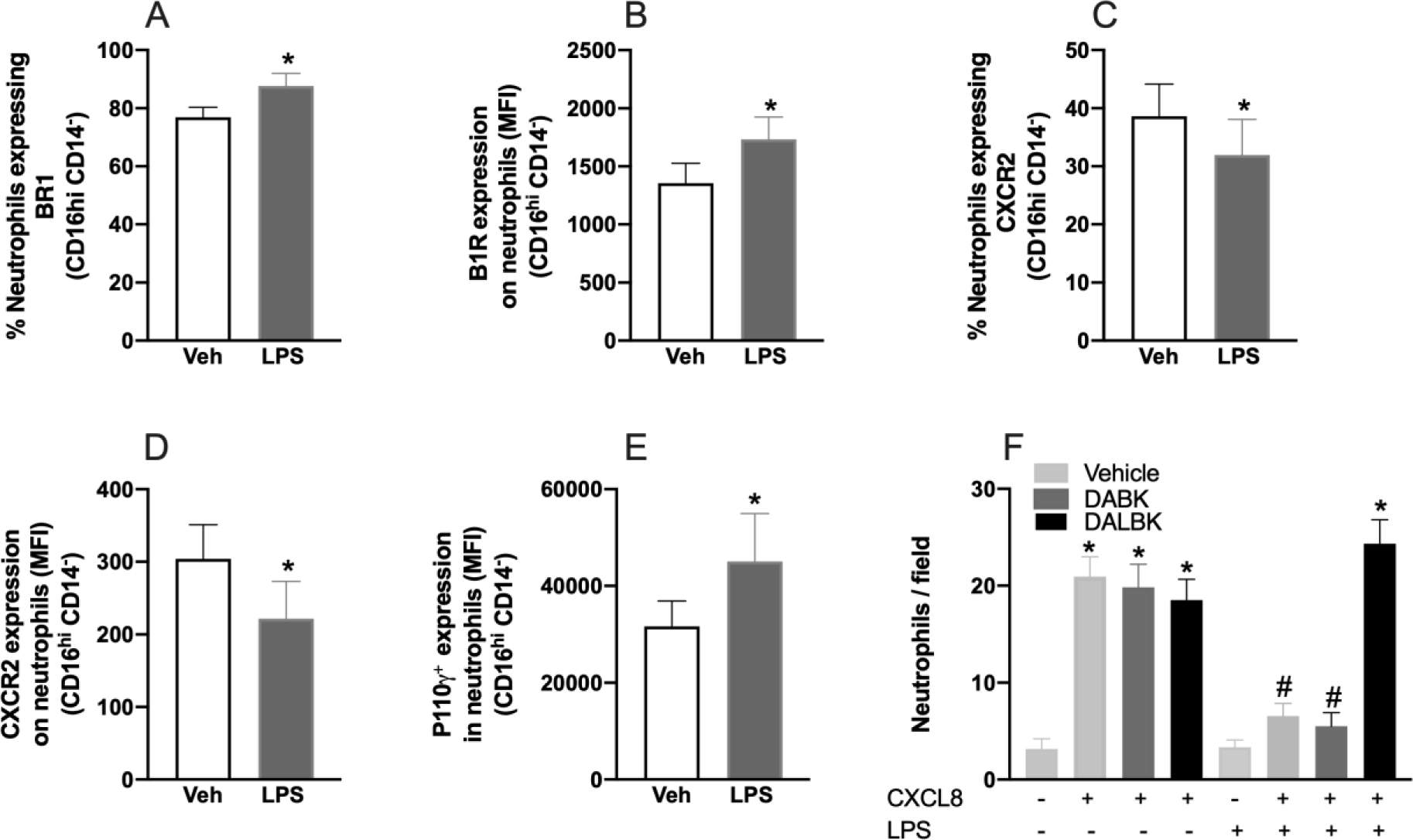
Inhibition of B1 in human neutrophils inhibits their ability to migrate under sepsis conditions. (A) Percentage of positive neutrophils and (B) mean fluorescence intensity for B1R, (C) percentage of positive neutrophils and (D) mean fluorescence intensity for CXRCR2, and (E) mean fluorescence intensity for p110 γ in neutrophils obtained from the blood of healthy individuals subjected to stimulation with LPS (10µM) or vehicle for two hours. Data correspond to mean ± EPM of 5 wells per group. * p<0.05 related to the vehicle group. (F) Chemotaxis of human neutrophils isolated from the blood of healthy individuals and those stimulated by LPS (10µM) for two hours. Migration was evaluated by Boyden chamber, using chemokine CXCL8 (10µM) as a chemotactic factor. B1 receptor agonist (Des-Arg9Bradykinin) (DABK-1 µM) and B1 receptor antagonist (Des-Arg9[Leu8]-Bradykinin) (DALBK-10µM) were incubated for 30 minutes with cells. In the last group, cells were incubated with B1 receptor agonist and antagonist simultaneously. Data correspond to mean ± EPM of four wells per group. * p<0.05 related to the control + group.

### 3.5. B1 antagonist improves sepsis and has a synergistic effect with broad-spectrum antibiotics

WT mice subjected to CLP were treated with DALBK, a B1 antagonist, 1 hour before (pre-treatment) or 6 hours after (post-treatment) the CLP procedure. Both pre-treatment and post-treatment with DALBK prevented the lack of migration of neutrophils into the peritoneal cavity (Fig. 6A, B), resulting in a reduction of bacterial load in the peritoneal lavage and serum (Fig. 6C) compared to vehicle treatment. Pre-and post-treatment with DALBK significantly reduced the TNF concentration in the peritoneal cavity, serum and lung (Fig.6D-F) induced by CLP. In addition, pre-or post-treatment with DALBK reduced CXCL-1 concentration in serum and lung (Fig.6H and I), although it had no effect on CXCL-1 concentration in the peritoneal cavity (Fig. 6G). B1 blockade also reduced CLP-induced MPO activity in the lung (Fig 6J).

**Figure 6:**
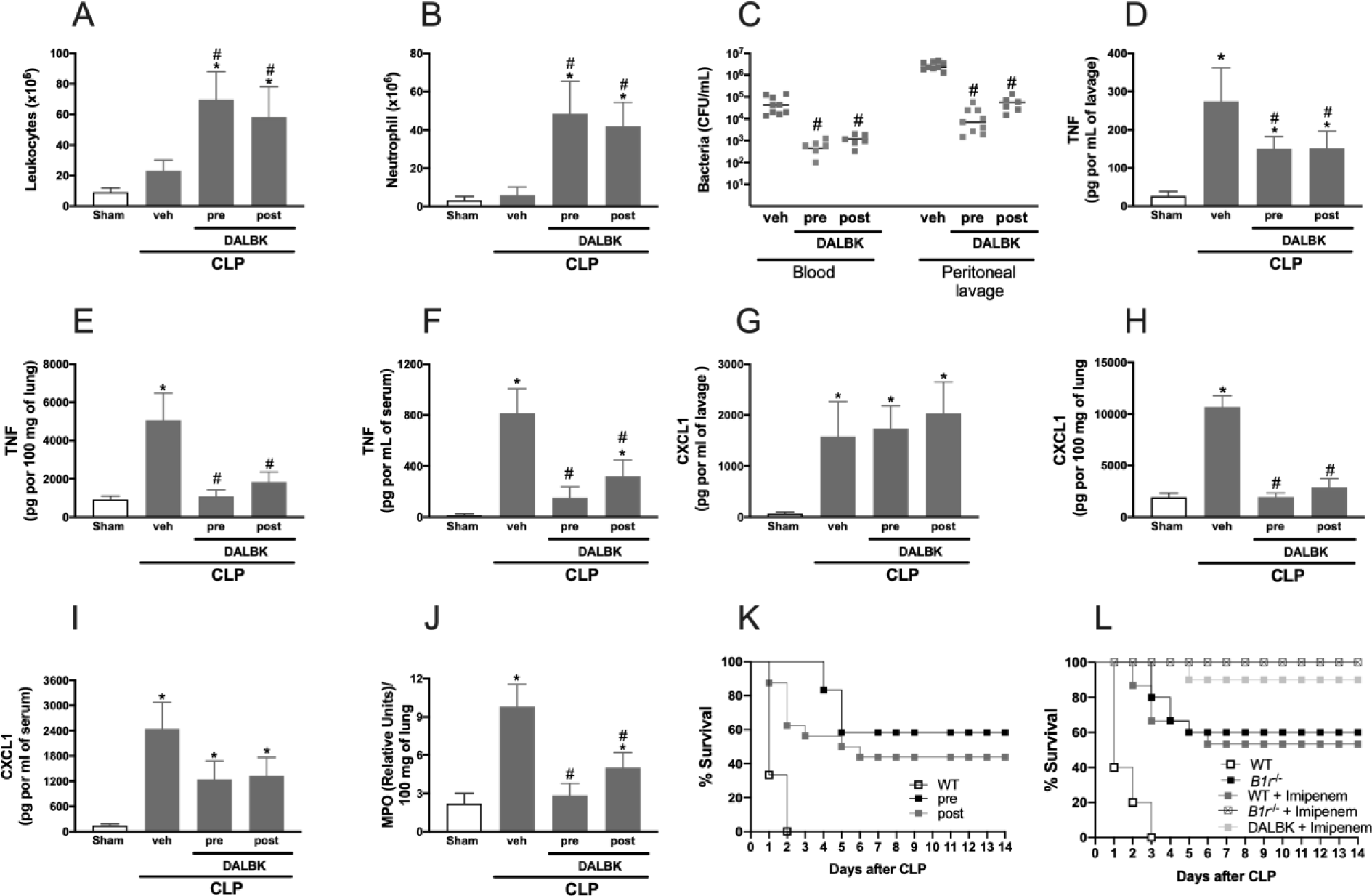
Treatment with a B1 antagonist prevents neutrophil failure, bacteria burden, inflammatory response, and lethality associated to CLP in WT mice. WT mice were treated with a B1 receptor antagonist (DALBK, 50nmolar/Kg) 1 hour before sepsis surgery (pre) or 3 hours after sepsis surgery (post). 6 hours after CLP, mice were euthanized blood and lungs were collected and peritoneal lavage was performed. (A) CFU, (B)total leukocytes, (C) neutrophil influx into the peritoneal cavity. The concentrations of (D) TNF and (E) CXCL1 were determined in the peritoneal lavage. The concentrations of (F) TNF and (G) CXCL1 were determined in the blood. (H) MPO activity, concentration of (I) TNF and (J) CXCL1 were measured in the lung. Results are expressed as mean ± SEM of at least five animals per group. *, *p* < 0.05, compared to the sham group and #, *p* < 0.05, compared to the wild-type animals submitted to CLP. (K) Survival of WT mice treated with a B1 antagonist or (L) the combined effect of post-treatment with a B1 antagonist and imipenem was followed for 14 days. Imipenem was administered 3 hours after sepsis surgery. Results are survival rates in %, analyzed with the Log-rank (Mantel-Cox) test, 10 mice per group. Results are expressed as mean ± SEM of at least five animals per group. *, *p* < 0.05, compared to the sham group and #, *p* < 0.05, compared to wild-type animals subjected to CLP.

Importantly, both pre- and post-treatment with DALBK reduced CLP-induced lethality. Approximately 50% of mice survived up to 14 days after sepsis (Fig. 6K). Since broad-spectrum antibiotics are the recommended treatment for septic patients, we next examined the effect of treatment with DALBK in mice previously treated with imipenem. Figure 6L shows that imipenem, as well as DALBK alone, reduced CLP-induced lethality by approximately 50%. Interestingly, post-treatment with DALBK provided 90% protection against lethality in CLP-mice that had previously been treated with an antibiotic. *B1^-/-^* mice treated with antibiotics survived 100%. Taken together, these results suggest that DALBK could be a potential agent for the treatment of sepsis.

## 4 Discussion

Despite numerous advances in the use of antimicrobials and resuscitation therapies for sepsis (17), the mortality rate remains high, and new therapeutic strategies for the treatment of this disease are urgently needed. This study shows that (i) B1 plays an essential role in the pathogenesis of sepsis, at least in part by mediating impaired neutrophil migration during the disease; (ii) B1 exerts its effect in myeloid cells by controlling the activation of P13ky and the expression of CXCR2; (iii) B1 antagonist improves the benefit of antibiotic treatment, highlighting DALBK as a potential adjuvant treatment for sepsis.

Previously, Ruiz et al. had showed that a B1 antagonist reduced lethality, lung and kidney injury, and vascular permeability in a CLP-induced septic shock model (18). Here, we demonstrated that B1 antagonism or its absence attenuates the severity of sepsis by improving neutrophil recruitment. Improved neutrophil migration facilitated bacterial clearance, reduced systemic cytokine production and mortality. BKB2R blockade had no effect on local inflammation, bacterial burden or mortality.

B1 expressed on neutrophils (19) can induce chemotaxis (20), enhance the expression of integrin (21), and trigger the release of matrix metalloproteinases (MMP)-9 and myeloperoxidase in a PI3K- and PKC-dependent manner in human neutrophils (22). Our study showed that the absence of B1 limits the overproduction of cytokines. Previous results show that the absence of B1 reduces TNF production in the intestine, lung and serum during intestinal ischemia and reperfusion in mice (6). Lower cytokine concentrations are associated with lower lethality. Under our experimental conditions, blocking B1 results in a more efficient, self-limiting inflammatory response that allows clearance of pathogens without excessive tissue damage. Our data are consistent with the findings of Nasseri and colleagues, who showed that the B1 antagonist BI113823 reduced the expression of NF-kB and COX-2 in the lungs of mice following LPS-induced lung injury, resulting in reduced lung injury (23). This study also showed that the same antagonist reduced cytokine concentrations and ameliorated lung injury in a polymicrobial sepsis rat model. Our results have shown that the role of B1 in sepsis appears to be related to its presence in bone marrow-derived cells, as demonstrated using chimeric mice.

Activation of B1 appears to be associated with internalization of CXCR2 in circulating neutrophils during severe sepsis. B1-deficient mice exhibited increased CXCR2 receptor surface expression. This was associated with decreased expression of P110γ, a subunit of phosphoinositide 3-kinase gamma (PI3Kγ), which is involved in the internalization of CXCR2 (24). Inhibition of PI3Kγ resulted in down-regulation of GRK2 expression, thereby reducing in CXCR2 internalization which contributed to increased CXCR2 expression in neutrophils and improved survival in septic mice (24). Our results therefore suggest that kinins acting on B1 trigger PI3Kγ activation leading to CXCR2 internalization. Alternatively, kinins that induce NO production could reduce neutrophil migration (23). NO triggers the activation of soluble guanylate cyclases (GCs) as well as the formation of cyclic-GMP and the phosphorylation of protein kinase G (PKG) (25). Inhibition of sGC and PKG during sepsis improved survival by increasing CXCR2 expression and neutrophil migration. These results shed light on the complex mechanisms that influence neutrophil responses in sepsis.

Preincubation of neutrophils with LPS induces expression of B1 and triggers internalization of CXCR2, a hallmark of sepsis. Our study shows that a B1 antagonist reverses impaired chemotaxis in human neutrophils exposed to LPS and CXCL8, suggesting B1 activation in humans, and providing a potential target for sepsis treatment. Furthermore, blocking B1 increases the efficacy of antibiotic therapy. Combining antibiotics with modulators of the immune response is a promising approach, but previous agents have shown limited results (24, 26). Treatment with anti-TNF or IL-1Ra have not had a positive effect on mortality in septic patients (27), suggesting that new candidates are urgently needed. Inhibition of B1 improves neutrophil migration and bacterial clearance, making it a valuable therapeutic candidate for the treatment of sepsis. In summary, our research highlights B1 as a potential treatment target for sepsis, offering improved modulation of the inflammatory response and synergy with antibiotics.

## Data Availability statement

The data that support the findings of this study are available from the corresponding author DGS request.

## Acknowledgments

We are grateful for the financial support provided by Instituto Nacional de Ciência e Tecnologia (INCT), Coordenação de Aperfeiçoamento de Pessoal de Nível Superior (CAPES) and Conselho Nacional de Desenvolvimento Científico e Tecnológico (CNPq).

## Author contributions

Resende L, Teixeira MM and Souza DG created the study design; Resende L, Arifa RDN, Resende C, Silva MEF, Resende B, Tavares L and Reis A performed data acquisition. Resende L, Teixeira MM, Souza DG analyzed and interpreted data and performed statistical analysis; Resende C, Arifa RDN, Amaral FA, Teixeira MM, Fagundes CT and Souza DG prepared the manuscript.

## Permission to reproduce

The authors provide full permission to reproduce content of the article where relevant.

---

**Table S1.**
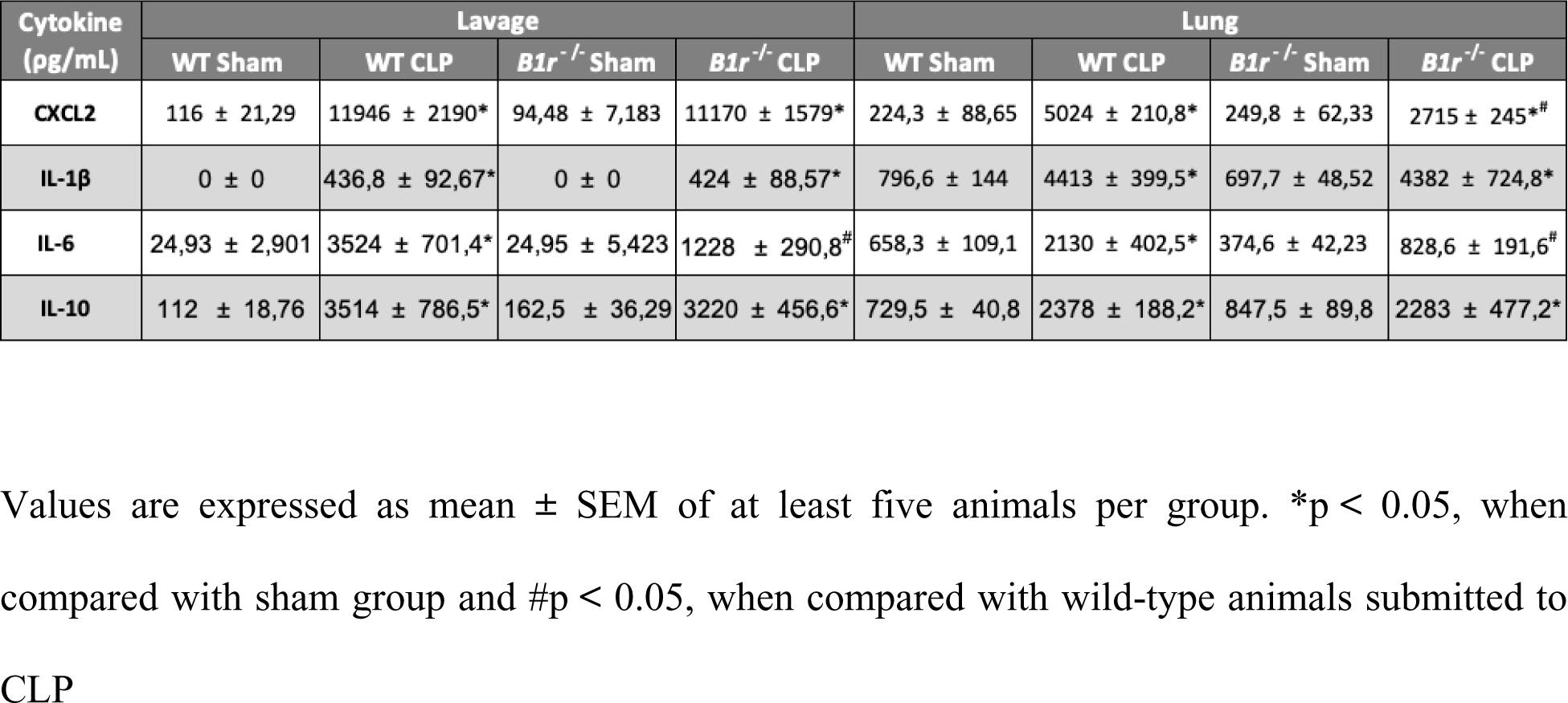
Cytokine production in the lavage fluid and lung tissues of both WT and *B1r*^-/-^ mice subjected to cecal ligation and puncture (CLP).

